# Linking individual differences in cognitive control and perception: Meta-perception in older and younger adults

**DOI:** 10.1101/2021.07.07.451419

**Authors:** Lena Klever, Pascal Mamassian, Jutta Billino

**Affiliations:** Experimental Psychology, Justus Liebig University Giessen, Germany; Center for Mind, Brain, and Behavior (CMBB), University of Marburg and Justus Liebig University Giessen, Germany; Laboratoire des Systèmes Perceptifs (CNRS UMR 8248), Département d’Études Cognitives, École Normale Supérieure, PSL University, 75005, Paris, France

## Abstract

Visual perception is not only shaped by sensitivity, but also by confidence, i.e. the ability to estimate the accuracy of a visual decision. There is robust evidence that younger observers have access to a reliable measure of their own uncertainty when making visual decisions. This metacognitive ability might be challenged during aging due to increasing sensory noise and decreasing cognitive control resources. We investigated age effects on visual confidence using a confidence forced-choice paradigm. We determined discrimination thresholds for trials in which perceptual judgements were indicated as confident and for those in which they were declined as confident. Younger adults (19-38 years) showed significantly lower discrimination thresholds than older adults (60-78 years). In both age groups, perceptual performance was linked to confidence judgements, but overall results suggest reduced confidence efficiency in older adults. However, we observed substantial variability of confidence effects across all particpants. This variability was closely linked to individual differences in cognitive control capacities, i.e. executive function. Our findings provide evidence for age-related differences in meta-perceptual efficiency that present a specific challenge to perceptual performance in old age. We propose that these age effects are primarily mediated by cognitive control resources, supporting their crucial role for metacognitive efficiency.

## Introduction

Human behaviour and its underlying neural mechanisms are mostly studied with a specific focus on a particular functional domain, e.g., perception, cognition, motivation or motor functions. Although this approach has allowed detailed models and theories, complexity of behaviour can only be captured comprehensively when also interactions across behavioural domains are considered (Basso & Suzuki, 2017; Braver et al., 2014; O’Callaghan et al., 2016). A particularly influential, well-investigated higher-level concept that shapes behaviour is metacognition. It refers to the ability to evaluate the quality and consequences of one’s own thoughts and behaviours (Flavell, 1979; Fleming & Frith, 2014). Metacognitive abilities are thought to be key for optimizing performance by balancing actual outcome and subjective estimates of its quality. While individual differences in behavioural performance are immediately obvious, the role of metacognitive resources has been rarely explored.

Ageing provides a powerful proxy to individual differences and offers a unique window to possible variability in metacognitive efficiency. Ageing, from a behavioural perspective, can be understood as an umbrella term that incorporates gradually changing resources in all functional domains and at the same time adaptive mechanisms that can stabilize performance. Although the view of aging as a process of deterioration and decline might be still prominent, understanding of age-related differences has gradually shifted towards a more complex characterization, including stability, decline, and compensation (e.g., Billino & Pilz, 2019; Cabeza et al., 2018; Hartshorne & Germine, 2015). Metacognition could crucially contribute to optimizing performance in the face of age-related resource decline (see Owsley, 2011; Park & Reuter-Lorenz, 2009; Samanez-Larkin & Knutson, 2015; Seidler et al., 2010). However, evidence so far has remained ambiguous.

Since the prefrontal cortex has been consistently identified as critical neural functional correlate of metacognition (Fleming, Huijgen, & Dolan, 2012; Morales et al., 2018; Valk et al., 2016), vulnerabilities during aging have been assumed. Prefrontal areas are subject to the most pronounced age-related volume loss (Kennedy et al., 2009; Walhovd et al., 2011). In addition, consistent with the involvement of prefrontal cortex, metacognition is considered as closely related to general higher-order cognitive processes, i.e., executive function (e.g., Fernandez-Duque et al., 2000; Roebers, 2017). Age-related decline in executive function, indeed, is the most prominent facet of cognitive ageing (Hasher & Zacks, 1988; Lacreuse et al., 2020; West, 1996). However, notwithstanding these clear predictions, it seems still a matter of debate how sensitive metacognition is to age.

The majority of studies that have investigated age-related differences in metacognition so far has focused on memory performance in older and younger adults, so called meta-memory (cf., Palmer et al., 2014). Meta-memory is typically assessed by subjective measures of how confident individuals feel about the quality of their own memory performance, e.g., by giving a prospective or retrospective judgement on a rating scale. Several studies have reported an increased mismatch between actual performance and the judgements on one own’s abilities in older adults (Cauvin et al., 2019; Dodson et al., 2007; Hansson et al., 2008; Pansky et al., 2009; Perrotin et al., 2006; Soderstrom et al., 2012; Toth et al., 2011; Wong et al., 2012). Older adults tend to be overconfident about the quality of their memory performance. On the other hand, there are almost as many studies that have found only minor or even no age effects on the accuracy of meta-memory (Hertzog & Touron, 2011; Lachman et al., 1979; Palmer et al., 2014; Voskuilen et al., 2018). Metacognition in other functional domains, e.g., problem solving, linguistics, perception, even seems to elude any age effects (Filippi et al., 2020; Geurten & Lemaire, 2019; Palmer et al., 2014). Heterogenous results might be due to the use of rating scales for assessing confidence. Ratings could be confounded with individual biases how to anchor judgements so that evaluation of metacognition is complicated (see Mamassian, 2016; Morgan et al., 1997). Moreover, confidence judgements in commonly used cognitive tasks are made on rather complex decisions involving multiple criteria that might underlie additional biases that are hard to control.

Given these issues, the investigation of metacognition in perceptual tasks, i.e., meta-perception, has attracted increasing consideration over the last years (for review, see Mamassian, 2016). Perceptual tasks qualify for a well-structured assessment of metacognition since they typically are characterized by simple decisions on given sensory evidence, e.g., contrast or orientation discrimination. These decisions are accompanied by a subjective sense of (un)certainty, depending on the strength of sensory signals. Having access to a reliable measure of one’s own uncertainty is a crucial aspect of perceptual confidence. Confidence in the correctness of one own’s decisions is basically correlated with accuracy of decisions (e.g., Fleming, Dolan, & Frith, 2012; see also Peirce & Jastrow, 1884). Observers will report high confidence when their perceptual decision is objectively correct, and low confidence when it is objectively incorrect. During ageing the quality of confidence judgements in perceptual tasks might be challenged in particular by pronounced age-related sensory decline due to peripheral vulnerabilities and increasing noise in neural representations (Fu et al., 2013; Owsley, 2011; Yang et al., 2009; Yu et al., 2006). However, only a single study so far has considered age effects on meta-perception. Palmer and colleagues (2014) investigated confidence ratings on contrast-defined pop-out detection in a sample covering the adult age range. They used meta d-prime as measure of metacognitive performance (Maniscalco & Lau, 2012) and determined decreased efficiency with increasing age. Findings though remain incoherent since the same sample showed no age-related decline in metacognition in a memory task using the same rating procedure. In addition, metacognitive efficiency in the perceptual as well as in the memory task was dissociated from executive function.

We aimed to scrutinize how age affects meta-perception using a confidence forced-choice paradigm (Barthelmé & Mamassian, 2009, 2010; see also Mamassian, 2016). This method has been proposed to derive a bias-free measure of confidence and avoids confounds emerging from confidence rating scales. Confidence measures in this paradigm are not affected by possible idiosyncratic confidence biases that have been reported in older adults (e.g., Cauvin et al., 2019; Hansson et al., 2008). It allows analyses based one the signal detection framework, controlling for differences in task performance. We hypothesize that older adults show decreased meta-perceptual efficiency and these age effects are crucially driven by individual differences in cognitive control capacities, i.e. executive function.

## Methods

### Participants

A total of 30 younger adults (18 females) and 30 older adults (17 females) participated in this study. The participants’ age ranged from 19 to 38 years with a mean of 24.6 years (*SD* = 4.4) in the younger group and from 60 to 78 years with a mean of 68.8 years (*SD* = 4.7) in the older group. Recruitment of participants was managed by calls for participation at the University of Giessen and in local newspapers. Any history of ophthalmologic, neurologic, or psychiatric disorders as well as medications presumed to interfere with visual functioning were screened out by a detailed interview protocol. Visual acuity was measured binocularly, confirming normal or corrected-to-normal for all participants. Older adults were screened for mild cognitive impairment using a cut-off score of ≥ 26 on the Montreal Cognitive Assessment Scale (Nasreddine et al., 2005). Methods and procedures were approved by the local ethics committee at Justus Liebig University Giessen and adhered to the principles of the Declaration of Helsinki (World Medical Association, 2013). Participants were compensated with course credits or money.

### Assessment of individual differences in cognitive abilities

We characterized cognitive abilities of our participants using a battery of established measures that particularly allowed for evaluation of executive function (EF). Critical measures were chosen in order to cover key facets of cognitive control processes (Miyake & Friedman, 2012) and included: the Digit Symbol Substitution Test (DSST) (Wechsler, 2008) measuring updating ability; the Trail Making Test part B (TMT-B) (Reitan & Wolfson, 1985) measuring shifting ability; the Victoria Stroop Test colour naming (VST-C) (Mueller & Piper, 2014; Stroop, 1935) measuring inhibition ability; the LPS-3 (Kreuzpointner et al., 2013), a subtest of a major German intelligence test battery, measuring nonverbal reasoning ability. These measures were combined into a global EF score for each participant by averaging the *z*-scores obtained for the individual measures. In addition, we assessed the maximal backward digit span (Härting et al., 2000) in order to evaluate short-term memory capacity that qualified as a possible confounding issue given the procedural details of our meta-perceptual task.

### Setup and stimuli

Visual stimuli were presented on a calibrated 32-inch Display++ LCD monitor (Cambridge Research Systems, Rochester, UK) with a spatial resolution of 1920 × 1080 pixels and a refresh rate of 120 Hz noninterlaced. The setup was placed in a darkened room and participants were seated at a distance of 100 cm in front of the monitor, resulting in a display size of 41° × 23°. White and black pixels had a luminance of 112.7 and 0.1 cd/m^2^, respectively, measured with a CS-2000 Spectroradiometer (Konica Minolta). Stimulus presentation was controlled by MATLAB using the Psychophysics toolbox (Brainard, 1997; Kleiner, 2010). A standard gamepad was used as input device (Microsoft SideWinder).

Stimuli were vertical Gabor patches displayed on an average grey background. Sinusoidal gratings had a spatial frequency of 0.8 cyc/°with randomized phase and the standard deviation of the Gaussian envelope was 1°. The contrast of the Gabor patches was sampled from seven different levels ranging from 13% to 31% in steps of 3%. The stimulus configuration consisted of two Gabor patches presented to the left and right of a central fixation dot at 4.2° eccentricity along the horizontal meridian. The fixation dot was black and had a diameter of 0.2°. One Gabor patch, i.e. the standard patch, had a fixed contrast of 22%, whereas the contrast of the other Gabor patch, i.e. the test patch, varied. Laterality of standard and test patches, respectively, was randomized.

### Procedure

We assessed meta-perception using a confidence forced-choice paradigm (Barthelmé & Mamassian, 2009, 2010). Figure 1 depicts a typical trial.

**Figure 1.**
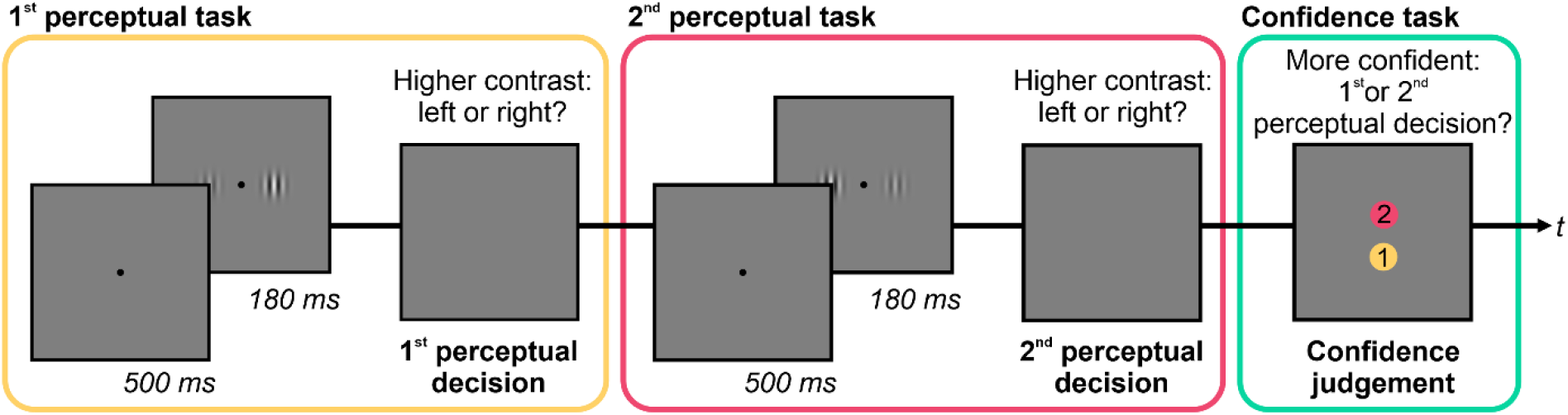
Trial procedure of the confidence forced-choice paradigm. Participants were presented with two consecutive perceptual tasks in which they had to decide which of two simultaneously presented Gabor patches appeared higher in contrast. After the second perceptual decision, they were asked for a confidence judgement, i.e. they had to indicate about which of the two perceptual decisions they felt more confident. Please note that colour is here used to illustrate the consecutive steps in each trial and was not used in the actual procedure.

Each trial consisted of two consecutive perceptual tasks, specifically contrast discrimination tasks, and a final confidence task. A fixation dot was shown for 500 ms, which was followed by two Gabor patches presented simultaneously for 180 ms. Then the display turned grey and participants decided whether the left or right patch appeared higher in contrast (first perceptual decision). Responses were entered with the respective index fingers using the trigger buttons on the back of the gamepad. Then an equivalent second task followed using different patches and another contrast decision was made (second perceptual decision). Afterwards, participants indicated about which of the two perceptual decisions they felt more confident (confidence judgement). The response was given with the right thumb using two vertically aligned buttons on the top side of the gamepad. The buttons were mapped to the first or second perceptual decision, respectively. The mapping was visualized on the display and balanced across participants.

Before data collection, a detailed instruction protocol and sufficient practice trials backed up that participants were familiar with the stimulus configuration, could comfortably follow the trial procedure, and handled the gamepad effortlessly. Following, participants completed a total of 420 trials, subdivided into 6 blocks with 70 trials each. Contrast levels of the test patches in the two consecutive contrast discrimination tasks were separately varied according to the method of constant stimuli, i.e. each of the 7 contrast levels was presented in 60 trials for the first and second contrast discrimination task, respectively.

### Data analyses

Based on participants’ confidence judgements, we divided perceptual decisions into two confidence sets: The first set included perceptual decisions that were chosen in the confidence task, i.e. they were associated with a relatively higher confidence, and was therefore labelled as *chosen*. The second set considered the ensemble of all perceptual decisions and was labelled as *unsorted*. We analysed perceptual performance for both sets by fitting cumulative Gaussian functions to the percentage of responses in which observers reported the contrast of the test patch as higher than the standard patch. The inverse standard deviation of these functions yields the contrast sensitivity. We used the psignifit 4 toolbox in Matlab that provides an accurate Bayesian estimation of psychometric functions and has been shown to be robust to overdispersion in measured data (Schütt et al., 2016). Goodness of fit of the psychometric functions was assessed with the measure of deviance *D* which supported good fits between the model and the data. Both sets showed similar Goodness of fit measures (*t*(59) = 1.82, *p* = .074, 95% CI [-0.117, 2.506], *d* = 0.26).

We quantified meta-perceptual sensitivity, i.e. the ability to estimate the accuracy of a perceptual decision, by a confidence modulation index (CMI) according to Equation 1. The CMI gives the sensitivity increase for the set of decisions chosen as confident relative to the set of unsorted decisions as a percentage of the sensitivity derived from the unsorted decisions.

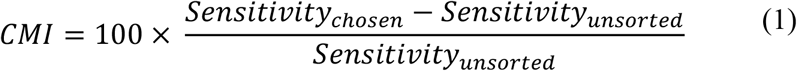

An individual observer who derives her confidence judgements completely dissociated from her perceptual decisions will show a CMI close to zero. However, the closer the confidence judgement is linked to the actual accuracy of the perceptual decision the higher the CMI will be, indicating better meta-perceptual sensitivity. Given that the CMI provides a proportional measure, values were arcsine-square-root transformed before they were submitted to statistical procedures. Inspecting the distribution of CMIs in our sample, we identified outlier data for one older participant. Her CMI deviated more than 1.5 times the interquartile range from the range borders of the complete sample. In order to enhance validity of our data and reduce unsystematic noise we discarded this participant from our analyses.

Time measures for perceptual decisions were explored using median response times (RT). Since perceptual decision times vary with stimulus difficulty and confidence in a given task, we disentangled both parameters by an elaborate analysis described in detail in previous studies (De Gardelle et al., 2016; De Gardelle & Mamassian, 2014). We first normalized stimulus values for each individual considering their psychometric functions. We calculated the signed distances *S* between the 7 used stimulus intensities and the point of subjective equality in standard deviation units of the psychometric function. Chosen and unsorted confidence sets were considered separately. We then fitted an exponential model with three free parameters to the median RTs for each of the 7 stimulus intensity levels. The model is defined by Equation 2. *RT(S)* gives the fitted RT for a normalized stimulus intensity level *S. C* gives the according mean confidence across all included perceptual decisions. We encoded confidence with 1 for perceptual decisions which were selected in the confidence choice task and with 0 for perceptual decisions which were not chosen.

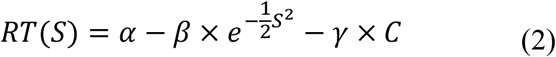

The model yields three parameters, i.e. α, giving the generic RT, β, capturing the exponential change in RT due to differences in stimulus intensity, and γ, capturing the linear change in RT due to confidence.

Sensitivity and RT data were analysed by mixed ANOVAs with the within-subject factor *condfidence set* (chosen vs. unsorted) and the between subject factor *age group* (older adults vs. younger adults). *T*-tests were used for age group comparisons of the CMI, cognitive measures, and RT parameters. If Levene’s test indicated unequal variances, degrees of freedom were adjusted appropriately. Associations between CMI and critical parameters investigated by correlational analyses. For group comparisons and correlational analyses, we computed 95% percentile confidence intervals using 2000 bootstrap samples. A significance level of α = .05 was applied for all statistical analyses and tests were two-sided. If not stated otherwise, descriptive values are given as means ± 1 SEM.

## Results

We initially explored the overall response patterns of older and younger adults in the confidence forced-choice paradigm. Age effects on meta-perceptual abilities were then analysed in detail by exploiting contrast sensitivity functions derived from the chosen und unsorted confidence sets, respectively. Differences in meta-perceptual sensitivity were scrutinized considering the role of processing speed and executive functions.

### Overview of response pattern

Figure 2 illustrates confidence judgements for perceptual decisions at different task difficulty levels, i.e. different contrast differences between the standard and test Gabor patches. The separation of data for correct and incorrect decisions provides a rough overview of meta-perception in our paradigm.

**Figure 2.**
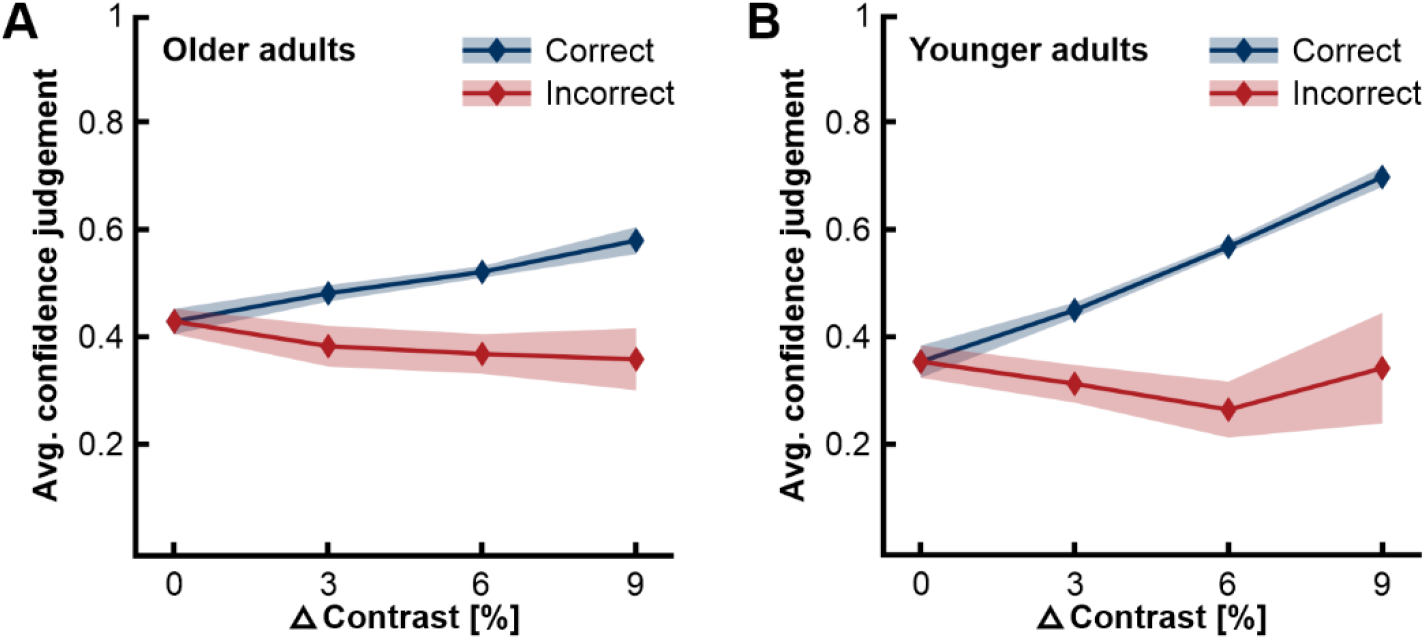
Average confidence judgements for perceptual decisions at different task difficulty levels, plotted separately for correct and incorrect decisions. (A) Data for older adults. (B) Data for younger adults. Task difficulty level is given as absolute contrast difference between the standard and test patches. Please note that task difficulty decreases with difference values. Confidence judgements were coded as 1 for chosen and as 0 for not chosen. Shaded areas give 95% confidence intervals.

In general, participants more often felt confident about their perceptual decisions if these were correct than if these were incorrect, indicating that they evaluated their performance appropriately. This difference in average confidence judgements for correct and incorrect decisions increased with decreasing task difficulty. Data pattern hence support that our paradigm captured meta-perceptual abilities in both age groups. However, Figure 2 also suggests age-related differences since the separation of data for correct and incorrect decisions is clearly less pronounced in older adults.

An elaborated illustration of the confidence judgement pattern in older and younger adults is given in Figure 3. Probabilities of confidence judgements are mapped with regard to task difficulties as well as correctness of perceptual decisions. The probabilities of choosing the first perceptual decision as more confident are mapped onto a coordinate plane defined by the stimulus strengths given in the first and second perceptual task, respectively. Separate maps for each of the four possible combinations of consecutive decisions are provided. The ability to judge the quality of the perceptual decisions is visualized in each map by a pattern of probabilities that dynamically varies in two dimensions.

**Figure 3.**
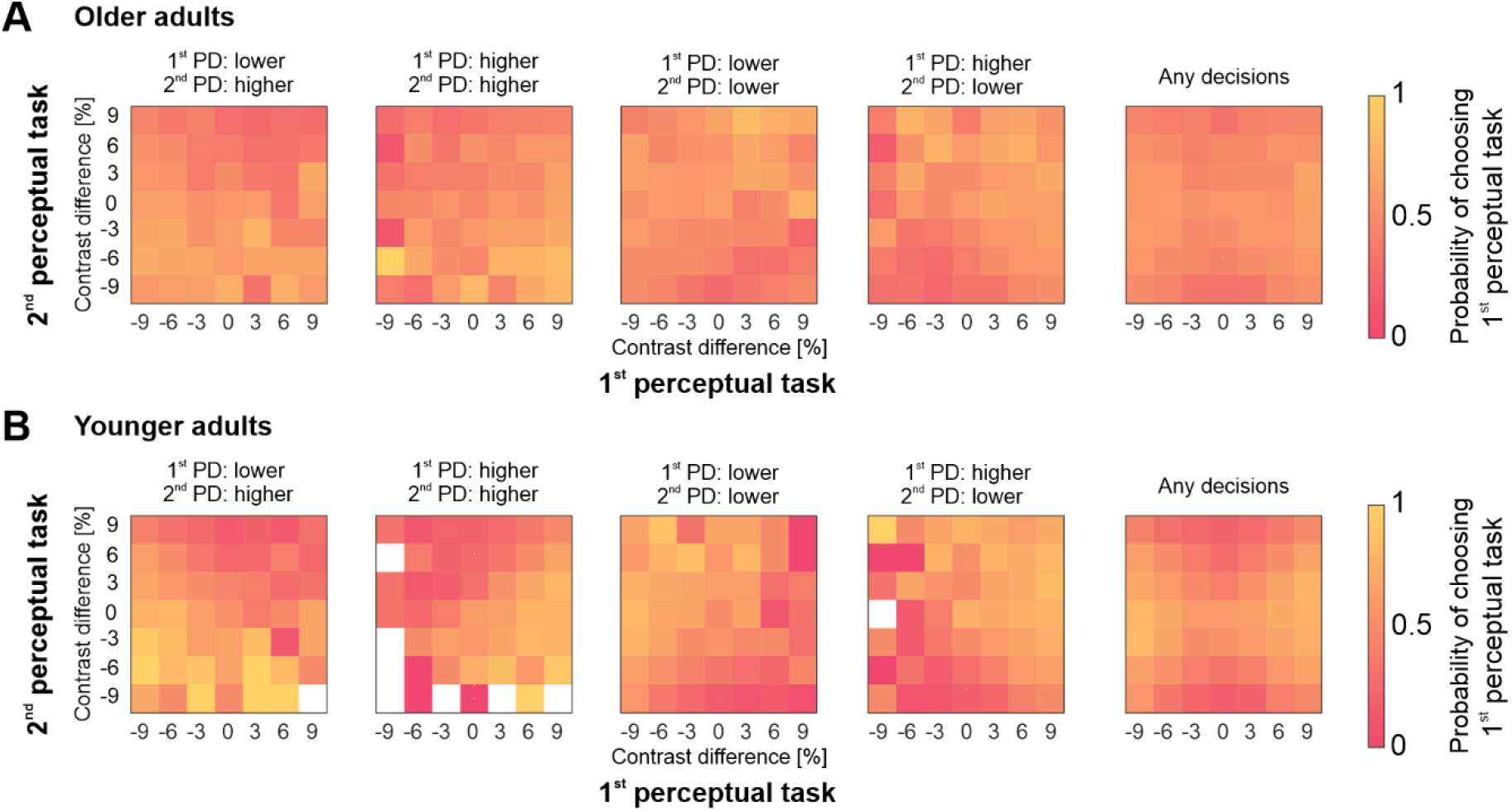
Descriptive illustration of meta-perceptual abilities in (A) older adults and (B) younger adults. The first four plots in each panel show the probability of choosing the 1st perceptual decision (PD) in the confidence judgement, i.e. associating it with relatively higher confidence, for each of the four possible combinations of perceptual decisions in the two consecutive contrast discrimination tasks. Decisions here apply to the test patches, i.e. code whether the test patches were indicated as lower or higher in contrast. The last plots on the right show the probability across all trials. The *x*- and *y*-axes give the contrast difference between the test patches and the standard patch in the first and second perceptual tasks, respectively. Meta-perceptual ability is indicated in these plots by a pattern of probabilities that dynamically depends on task difficulty, i.e. absolute contrast difference, and correctness of the perceptual decisions in both consecutive tasks. Please note that in white areas probabilities are not defined since the specific combination of consecutive perceptual decisions and stimulus strengths did not occur in our data set.

Generally, probabilities should gradually increase with contrast difference values in the first task that are consistent with the first perceptual decision. In parallel, they should gradually decrease with contrast difference values in the second task that are consistent with the second perceptual decision. Based on these dynamics specific triangular patterns emerge for each of the decision combinations. Better meta-perceptual abilities are reflected by more pronounced triangular patterns. Finally, mapping probabilities pooled across all possible combinations of perceptual decisions allows evaluation of possible choice preferences linked to the task order. A symmetric pattern of probabilities anchored at minimal stimulus strengths indicates the absence of systematic biases. Confidence probability maps for both older and younger adults suggest meta-perceptual abilities that allow for an overall appropriate evaluation of perceptual decisions. For neither age group we found evidence for task order biases that could confound confidence judgements. At the same time, the comparison between the maps for each age group again points to relevant age-related differences. Critical triangular probability patterns are prominent in younger adults, whereas in older adults the gradient of probabilities appears substantially blurred.

In summary, the exploration of response pattern in the confidence forced-choice paradigm suggests that in both age groups participants appropriately derived confidence judgements on their perceptual decisions and thus demonstrated meta-perceptual abilities. However, evidence for age-related differences emerges and is followed up by quantifying how close confidence judgements are linked to perceptual decisions.

### Psychometric analyses

We were initially interested in determining whether contrast sensitivity varies systematically between the two confidence sets, i.e. chosen or unsorted sets, and between the groups of older and younger adults. We consistently observed higher contrast sensitivity for the chosen confidence set than for the unsorted confidence set. Figure 4 shows example psychometric functions of contrast discrimination for a representative older (A) and younger (B) adult, respectively. For both participants, the functions derived from the two confidence sets differ in slope, indicating higher contrast sensitivity for the chosen confidence set. Points of subjective equality lie close to each other.

**Figure 4.**
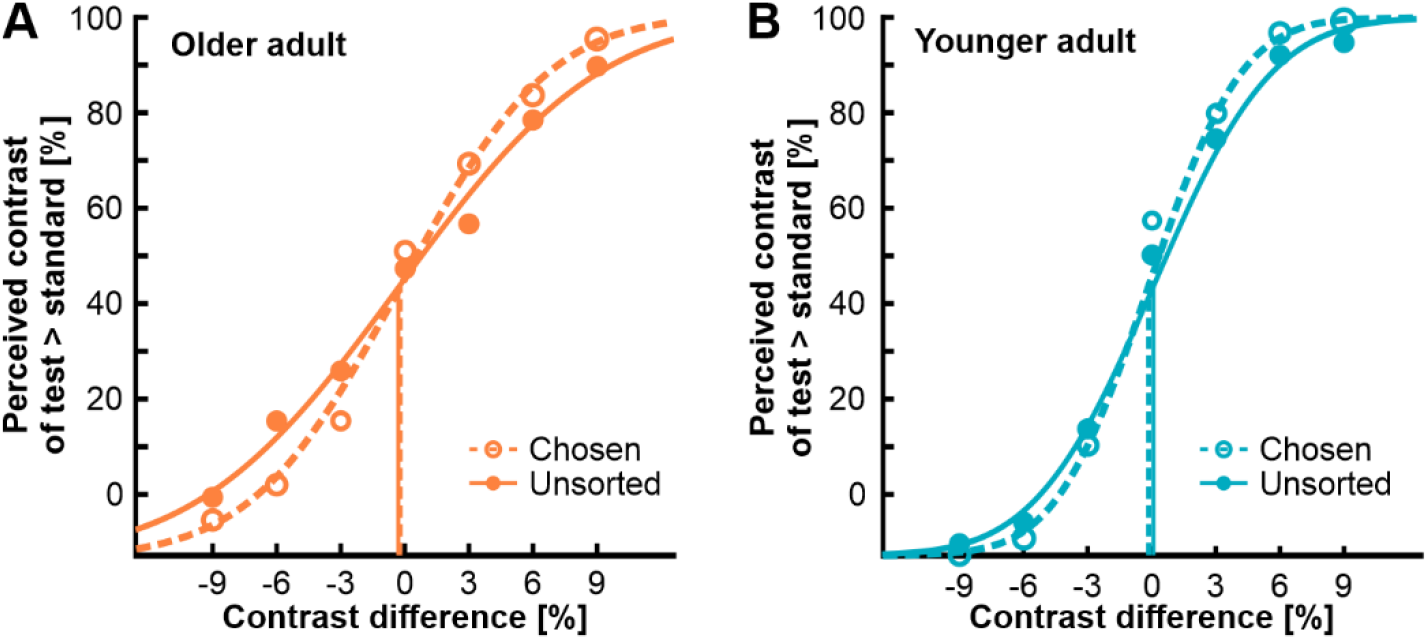
Psychometric functions of contrast discrimination for (A) an example older adult and (B) an example younger adult. Proportion of decisions indicating higher contrast of the test patch compared to the standard patch is plotted as function of stimulus intensity given as the contrast difference between the test patch and the standard patch. Dashed lines and open dots represent data from the chosen confidence set, solid lines and closed dots represent data from the unsorted confidence set.

Analysis of pooled sensitivity data corroborated inspection of the example psychometric functions. Figure 5A illustrates contrast sensitivity we determined for each confidence set in both age groups. We submitted sensitivity data to a two-factorial ANOVA with *age group* as between-subjects factor and repeated measures on the factor *confidence set*. The analysis yielded significant main effects of *age group*, *F*(1, 57) = 30.30, *p* < .001, η_p_^2^ = .35, and *confidence set*, *F*(1, 57) = 114.79, *p* < .001, η_p_^2^= .67. However, these main effects were qualified by a significant interaction between both factors, *F*(1, 57) = 14.67, *p* < .001, η_p_^2^= .21. The interaction effect was followed up by post-hoc *t*-tests. They corroborated lower sensitivities in older adults for both confidence sets (both *p*’s < .001). Effect sizes were similar, i.e. *d* = 1.39 for the chosen and *d* = 1.44 for the unsorted confidence set. The sensitivity advantage for the chosen confidence set was significant for both older and younger adults (both *p*’s < .001); however, the difference was less pronounced in older adults, i.e. *d* = 0.42 vs. *d* = 0.55, respectively.

**Figure 5.**
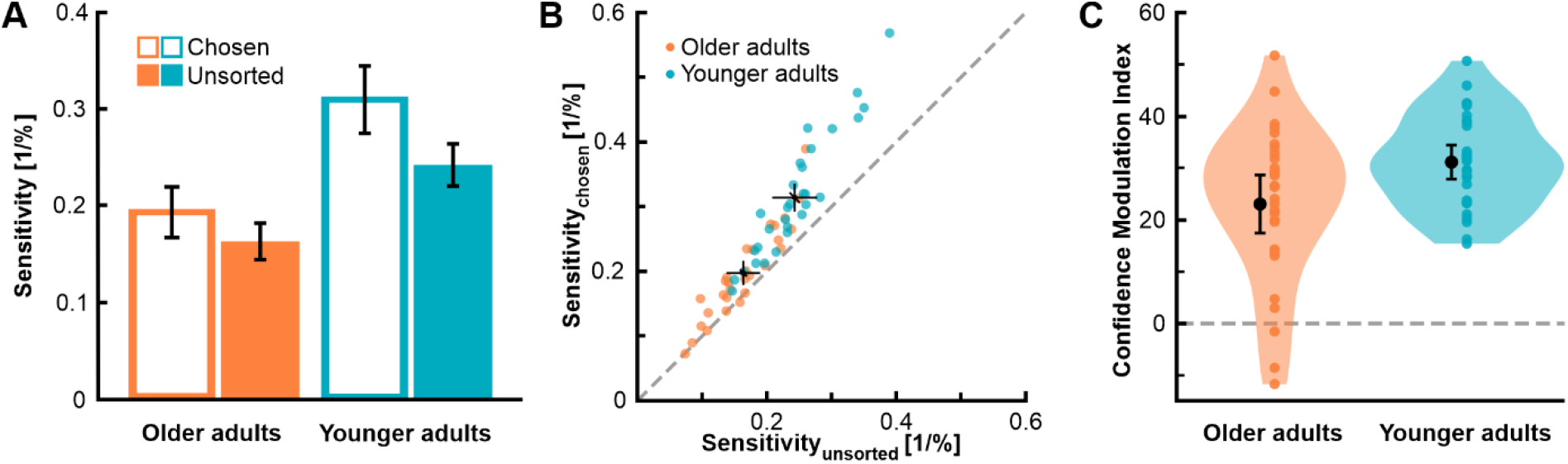
Contrast sensitivity and confidence. (A) Average contrast sensitivity as a function of age group and confidence set; open bars illustrate data from the chosen confidence set, closed bars represent data from the unsorted confidence set. (B) Contrast sensitivity for the chosen confidence set plotted against contrast sensitivity for the unsorted confidence set; each dot represents data from an individual participant, data for older and younger adults are plotted in different colours; dashed line marks the identity line; black closed dots give average sensitivities in each age group. (C) Confidence Modulation Index (CMI) as a function of age group; CMIs give the percental sensitivity increase from the set of unsorted trials to the set of chosen trials; coloured dots illustrate individual data and black dots represent the mean; shaded areas display 95% of the data distribution smoothed by a kernel density function. Error bars give 95% confidence intervals.

Figure 5B highlights these findings by giving a scatterplot of sensitivities for the unsorted confidence set against sensitivities for the chosen confidence set. Data for older and younger adults are illustrated in different colours. Whereas individual data points for younger adults lie exclusively above the diagonal identity line, those for older adults overall lie closer to and sometimes even marginally below it. Average values for both age groups show not only lower sensitivities, but also a smaller shift from the identity line in older adults. Confidence intervals suggest similar data precision in both age groups.

We further inspected whether the points of subjective equality (PSE) differ between the chosen and unsorted confidence sets. PSEs should logically lie close to zero, i.e. standard and the test patches should be indistinguishable when there is no contrast difference. A shift of PSEs for the chosen confidence set could indicate that confidence judgements rely on a biased criterion and thus meta-perceptual efficiency is inherently limited. Comparisons of PSEs for the chosen and unsorted confidence sets yielded consistent results. For older as well as for younger adults the PSEs for the chosen and unsorted confidence sets did not deviate from each other (older adults: *t*(28) = 0.06, *p* = .953, *d* < 0.01; younger adults: *t*(29) = −0.05, *p* = .960, *d* < 0.01).

### Meta-perceptual sensitivity

In order to investigate individual differences in meta-perceptual sensitivity, we analysed the sensitivity increase for the set of perceptual decisions chosen as confident relative to the set of unsorted decisions as a percentage of the sensitivity derived from the unsorted decisions, i.e. the CMI (see Methods). Figure 5C gives these meta-perceptual sensitivities for older and younger adults. We initially used one-sample *t*-tests to evaluate whether CMIs differed from zero. Results supported positive CMIs in the older age group, *t*(28) = 8.21, *p* < .001, *d* = 1.52, as well as in the younger age group, *t*(29) = 18.99, *p* < .001, *d* = 3.47. Participants in both age groups thus showed some ability to judge the validity of their perceptual decisions. However, on average, meta-perceptual sensitivity was lower in older, *M* = 23.04 ± 2.81, compared to younger adults, *M* = 31.21 ± 1.64. Whereas the link between confidence judgements and objective accuracy of perceptual decisions triggers a relative sensitivity benefit of over 30% in younger adults, the benefit is limited to less than 25% in older adults, *t*(45.34) = −2.51, *p* = .016, *d* = −0.66. Please note that we observed substantial variability of CMIs in our sample, especially pronounced in the group of older adults (Levene’s test: *F* = 4.87, *p* = .031). We next aimed to scrutinize which functional capacities drive the described age effect.

We were particularly interested in the role of cognitive control capacities since their decline essentially characterizes cognitive ageing. We captured cognitive control capacities by an EF score that covers key facets of this functional domain. Figure 6A gives EF scores in both age groups. On average, older adults showed less cognitive control capacities than younger adults, *t*(46.88) = −9.37,*p* < .001, *d* = −2.44.

**Figure 6.**
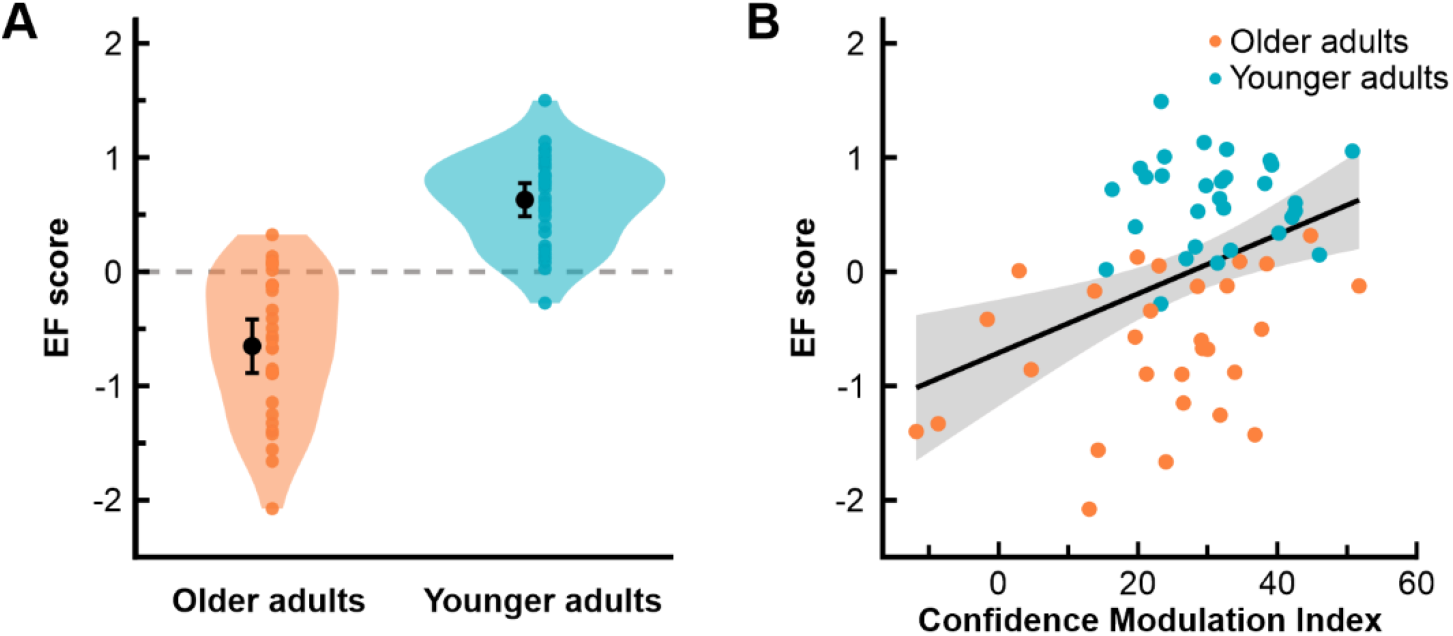
Cognitive control capacities and meta-perceptual sensitivity. (A) EF score as a function of age group; EF scores provide combined measure for cognitive control capacities averaging *z*-scores from DSST, TMT-B, VST-C, and LPS-3; coloured dots illustrate individual data and black dots represent the mean; shaded areas display 95% of the data distribution smoothed by a kernel density function. Error bars give 95% confidence intervals. (B) EF scores as a function of CMIs; data for older and younger adults are plotted in different colours; shaded area gives 95% confidence interval.

We investigated the link between meta-perceptual sensitivity and cognitive control capacities considering our complete sample in order to comprehensively exploit interindividual variability. Figure 6B illustrates the link between the CMI and the EF score. We determined a robust correlation of *r*(59) = .40, *p* = .001, 95% CI [.17, .57]. EF scores explained 16% of the variance in meta-perceptual sensitivity. Depiction of age group membership for each data point suggests that this correlation is not merely driven by group differences, but actually describes a general link. Consistently, a partial correlation analysis controlling for the factor age group, though attenuating the correlation, yielded corresponding results, *r*(56) = 0.26, *p* = .045. 95% CI [-.01, .50]. Our findings thus indicate that age-related differences in meta-perceptual sensitivity are crucially driven by cognitive control capacities.

Short-term memory capacity represents another cognitive resource that is subject to prominent age-related changes. Considering that the procedure of the confidence forced-choice paradigm putatively necessitates relevant memory resources, we wanted to check whether the age effect on meta-perceptual sensitivity can be explained by a confound inherent to the procedural task demands. The digit span measure we used to assess short-term memory capacity indicated significantly lower capacities in our older adult group, *t*(57) = −2.82, *p* = .007, *d* = −0.58. However, we found no evidence that the CMI is linked to individual differences in memory capacity, *r*(59) = .12, *p* = .363, 95% CI [-.17, .41]. Given this result, we consider it as rather unlikely that meta-perceptual sensitivity had been compromised by task demands that might be more challenging for older adults with lower memory resources.

We finally explored whether age-related slowing could contribute to differences in meta-perceptual sensitivity. Confidence efficiency might be costly in terms of processing time so that limited resources could hamper meta-perceptual sensitivity in older adults. First, we analysed median RTs by a two-factorial ANOVA with *age group* as between-subjects factor and repeated measures on the factor *confidence set*. Figure 7A shows average RTs as a function of age group and confidence set.

**Figure 7.**
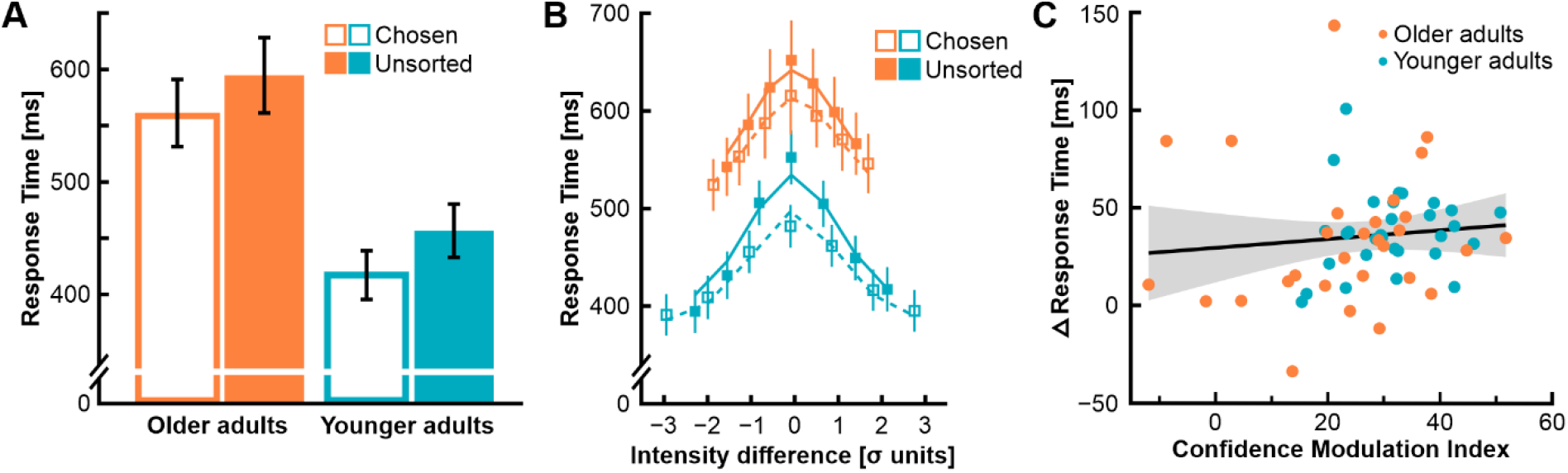
Response times (RT). (A) Average RTs as a function of age group and confidence set; open bars illustrate data from the chosen confidence set, closed bars represent data from the unsorted confidence set. (B) RTs for bins of stimulus intensities, i.e. contrast difference between test and standard patches, given in standard deviation units of the psychometric function (see Methods); squares represent average group data, lines represent the average fitted data; dashed lines and open squares represent data from the chosen trial set, solid lines and closed squares represent data from the unsorted trial set; colour code for age groups corresponds to (A). (C) RT differences between the unsorted and the chosen confidence sets as a function of CMIs; data for older and younger adults are plotted in different colours. Error bars and shaded areas give 95% confidence intervals.

We observed a significant main effect for *age group*, *F*(1, 57) = 13.38,*p* = .001, η_p_^2^ = .19, indicating slower RTs for older adults (chosen: *M* =561 ± 30 ms; unsorted: *M* =595 ± 33 ms) as opposed to younger adults (chosen: *M* =419 ± 21 ms; unsorted: *M* =457 ± 23 ms). In addition, a significant main effect of *confidence set* supported faster RTs for the chosen confidence set, *F*(1, 57) = 88.17, *p* < .001, η_p_^2^ = .61. There was no interaction between both main effects, *F*(1, 57) = 0.32, *p* = .572, η_p_^2^ < .01. The relationship between RTs and confidence turned out be similar in both age groups.

Since RTs are not only affected by confidence, but also by stimulus difficulty, we further aimed to clarify potential age-specific contributions. We disentangled both factors by modelling the RTs in each age group with three free parameters (see Methods). Fitting results are illustrated in Figure 7B. Consistent with the previous analysis, the first parameter α, giving the generic RT, significantly differed between the two age groups (older adults: *M* =524 ± 31 ms; younger adults: *M* =438 ± 25), corroborating age-related slowing, *t*(57) = 2.23, *p* = .030, *d* = 0.58. For both parameters β and γ, giving the influence of stimulus intensity and confidence on RTs, respectively, we determined values that consistently differed from zero for older and younger adults (all *p*’s < .001). RTs became slower with decreasing stimulus intensity, i.e. increasing difficulty, and faster with confidence. Most importantly, neither the parameter β nor the parameter γ differed between age groups (β: *t*(33.17) = −0.43, *p* = .672, *d* = −0.11; γ: *t*(57) = −0.71, *p* = .483, *d* = −0.18). These results corroborate that perceptual decision times underlie similar mechanisms in older and younger adults. Concluding, we directly tested whether the RT differences in the chosen relative to the unsorted confidence set were linked to meta-perceptual sensitivity. Figure 7C gives the RT differences as a function of the CMI. Both parameters were not significantly correlated, *r*(59) = .10, *p* = .450, 95% CI [-.18, .44]. Overall, RT analyses suggest that individual differences in meta-perceptual sensitivity do not emerge from processing speed dynamics.

## Discussion

Our perception relies on decisions about sensory evidence and the subjective confidence in the accuracy of these decisions. Visual perception is subject to massive age-related changes, however, the complexity of processes that contribute to these changes is still not well understood (see Billino & Pilz, 2019). In this study, we were concerned with age effects on meta-perception, i.e., the ability to judge the accuracy of one’s own perceptual decisions. Given age-related vulnerabilities in neural and cognitive resources that have been shown to be critical for metacognition, we supposed that meta-perceptual efficiency decreases with age.

We investigated meta-perceptual ability in a sample of healthy older and younger adults with an established confidence forced-choice paradigm that avoids idiosyncratic judgement biases (Barthelmé & Mamassian, 2009, 2010). We characterized participants’ executive function capacities using a comprehensive EF score that covers the key facets cognitive control. We were thus able to scrutinize the role of individual differences in cognitive control resources for meta-perceptual efficiency. Our results show that older adults still have access to a reliable measure of their uncertainty underlying perceptual decisions. Confidence judgements were consistently linked to the accuracy of perceptual decisions in both age groups. However, the efficiency of this link significantly decreases with age. While confidence judgements explained a sensitivity benefit of over 30% in younger adults, this benefit was limited to less than 25% in older adults. Across our participants we observed substantial individual differences in meta-perceptual sensitivity. We determined that 16% of variance in meta-perceptual sensitivity can be explained by individual cognitive control resources. Importantly, the critical impact of executive function was not exclusively defined by age-related differences, but showed as a general functional link that drives individual differences in meta-perception.

Our findings expand the understanding of how metacognition impacts perception across the adult lifespan. In our paradigm, we observed that older adults could selectively chose the interval that led to a higher performance in some cases. This indicates that they can evaluate the quality of their percepts. When compared to younger adults, though, this ability is reduced on average. This result is consistent with insights from the only previous study concerned with age effects on metacognition in perceptual tasks. Palmer and colleagues (2014) congruently reported reduced performance introspect with increasing age, using a contrast-defined pop-out detection task. Older adults showed lower awareness of their perceptual performance, but confidence was assessed by ratings scales which might have introduced confounds due to age-specific bias in confidence judgements (Cauvin et al., 2019; Hansson et al., 2008; see Mamassian, 2016; Morgan et al., 1997). The confidence force-choice paradigm avoids such biases so that our results corroborate independent meta-perceptual age effects.

Interpretation of our findings might be complicated by several factors that require careful consideration. Task difficulty might affect quality of confidence judgements. For our contrast discrimination task we chose sinusoidal gratings with a spatial frequency of 0.8 cyc/° for which age differences in contrast sensitivity were expected to be negligible (e.g., Owsley et al., 1983). We yet found clear age effects on contrast discrimination thresholds, putatively triggered by relatively short presentation times (cf., Bennett et al., 2007; Roudaia et al., 2011). Older adults showed higher thresholds and given that we used the method of constant stimuli for threshold measurement, higher task difficulty is implied for our group of older adults. Differences in task difficulty could, in turn, compromise confidence decisions (Maniscalco & Lau, 2012). For example, if the task is too difficult, an observer will not be good at identifying high confidence trials. Conversely, if the task is too easy, an observer might not be good at identifying low confidence trials. However, fit of the psychometric functions suggested that the applied intensity range was well-suited to capture performance across age groups. There was no difference between quality of fits in both age groups. We consider it as rather unlikely that probably unavoidable differences in task difficulty can explain the systematic age effects on the accuracy of confidence judgements. Furthermore, we ruled out that differential task difficulties emerging from short-term memory affordances explain age-related differences in meta-perception. Older and younger adults differed significantly in short-term memory resources, but we could not determine a relevant impact of this parameter on meta-perceptual sensitivity derived from our paradigm.

It might be also speculated whether differences in processing speed contribute to age effects on meta-perception. The reduction of processing speed is probably the most pronounced and robust functional age difference (Park & Reuter-Lorenz, 2009; Salthouse, 1996). Higher confidence in perceptual decisions is found to be associated with faster response times (De Gardelle et al., 2016; De Gardelle & Mamassian, 2014). This acceleration might be compromised and limited resources could hamper meta-perceptual sensitivity in older adults. As expected, we determined significantly prolonged response times in older adults compared to younger adults. However, response times were similarly modulated by confidence in both age groups. We found that independent of age responses were speeded up for perceptual decisions that are judged with higher confidence. In sum, we thus corroborate previous results showing differences in response times as a function of confidence in younger adults (De Gardelle et al., 2016; De Gardelle & Mamassian, 2014) and extend these findings to older age. Individual differences in processing speed do not interfere with efficient confidence judgements. In contrast, response times are consistently shaped by the confidence in the accuracy of perceptual decisions.

A main focus of our study was on the link between executive function and meta-perceptual sensitivity. Given the substantial conceptual overlap between metacognition, i.e. monitoring of decision quality, and executive function, i.e. cognitive control, a functional relationship suggests itself (Flavell, 1979; Fleming, Dolan, & Frith, 2012; Miyake & Friedman, 2012). In addition, both concepts have been shown to rely on shared neural resources (Fleming, Huijgen, & Dolan, 2012; Fuster, 2000; Morales et al., 2018; Valk et al., 2016). Ageing offers a powerful proxy to individual differences in executive function (Hasher & Zacks, 1988; Kennedy et al., 2009; Lacreuse et al., 2020; Park & Reuter-Lorenz, 2009; West, 1996). We captured individual cognitive control resources in a comprehensive score of executive function that was supposed to cover facets of the concept broadly (Miyake & Friedman, 2012). Older adults on average showed lower EF scores than younger adults, consistent with established findings on age effects on executive function (Park & Reuter-Lorenz, 2009). Thus, cognitive control resources could be identified a plausible candidate driver of age-related differences in meta-perceptual sensitivity. Most importantly, we were able to exploit the variability in EF scores across our older and younger participants to reveal a general functional link between cognitive control resources and meta-perception. This evidence is in line with several previous findings suggesting that metacognition basically relies on cognitive control resources (Maniscalco et al., 2017; Pansky et al., 2009; Souchay & Isingrini, 2004). We are aware of conflicting results indicating that metacognition and cognitive control might be better understood as independent capacities (Filippi et al., 2020; Palmer et al., 2014). However, we suggest that in some studies the functional links might be attenuated by executive function measures covering only specific facets of the concept. In addition, restriction of the range of individual differences in cognitive control resources due to very homogenous samples with regard to age and education can be assumed top obscure functional links.

Since our study was dedicated to meta-perception, it has to remain speculative whether our findings also hold for metacognition in other functional domains, e.g., meta-memory. Heterogeneity of results with regard to age effects on meta-memory hamper systematic evaluations (e.g., Dodson et al., 2007; Lachman et al., 1979; Pansky et al., 2009; Wong et al., 2012). Inconsistent results might primarily emerge from specific biases due to applied methods of measuring metacognitive parameters. At the same time there is evidence for general, domain-independent metacognitive mechanisms that suggests a coherent concept. (Maniscalco et al., 2017; McCurdy et al., 2013). We thus propose that our findings on age effects and the pivotal impact of cognitive control resources hold not only for meta-perception, but also for metacognitive mechanisms in other decision tasks.

To conclude, we showed that visual perception in older adults is shaped by metacognition. Older adults have access to a reliable measure of their own uncertainty when making visual decisions. Metacognitve capacities are key for behavioral control. For instance, a reduced performance introspect could result in not being able to identify relevant aspects of a task and inefficient allocation of resources (e.g., Desender et al., 2018). However, we found clear age-related differences in meta-perceptual sensitivity. Our results suggest reduced confidence efficiency in older adults. In principle, these age effects could be due to compromised reliability of judgements, but also due to declining cognitive control resources (cf., Bolenz et al., 2019). Exploiting individual differences across our complete sample, we corroborated the crucial functional role of cognitive control resources for metacognition. We propose that age effects on meta-perception are primarily mediated by this functional link. This finding is in line with converging evidence that age-related changes in perception and sensorimitmor control are critically driven executive contributions to efficient resource control (Chang et al., 2014; Huang et al., 2017; Huang et al., 2018; Monge & Madden, 2016).

## Acknowledgements

This work was funded by the German Research Foundation (Deutsche Forschungsgemeinschaft, DFG), Collaborative Research Centre SFB/TRR 135: Cardinal Mechanisms of Perception, project number 222641018. We thank Sabine Margolf for help with data collection. Data are publicly available at the doi: 10.xxxx/zenodo.xxxxxx

## Notes

### Competing Interest Statement

The authors have declared no competing interest.

## References

Barthelmé, S., & Mamassian, P. (2009). Evaluation of objective uncertainty in the visual system. PLoS Computational Biology, 5(9), e1000504. https://doi.org/10.1371/journal.pcbi.1000504

Barthelmé, S., & Mamassian, P. (2010). Flexible mechanisms underlie the evaluation of visual confidence. PNAS, 107(48), 20834–20839. https://doi.org/10.1073/pnas.1007704107

Basso, J. C., & Suzuki, W. A. (2017). The Effects of Acute Exercise on Mood, Cognition, Neurophysiology, and Neurochemical Pathways: A Review. Brain Plasticity, 2(2), 127–152. https://doi.org/10.3233/BPL-160040

Bennett, P. J., Sekuler, R., & Sekuler, A. B. (2007). The effects of aging on motion detection and direction identification. Vision Research, 47(6), 799–809. https://doi.org/10.1016/j.visres.2007.01.001

Billino, J., & Pilz, K. S. (2019). Motion perception as a model for perceptual aging. Journal of Vision, 19(4), 3. https://doi.org/10.1167/19.4.3

Bolenz, F., Kool, W., Reiter, A. M., & Eppinger, B. (2019). Metacontrol of decision-making strategies in human aging. ELife, 8. https://doi.org/10.7554/eLife.49154

Brainard, D. H. (1997). The Psychophysics Toolbox. Spatial Vision, 10(4), 433–436. https://doi.org/10.1163/156856897X00357

Braver, T. S., Krug, M. K., Chiew, K. S., Kool, W., Westbrook, J. A., Clement, N. J., Adcock, R. A., Barch, D. M., Botvinick, M. M., Carver, C. S., Cools, R., Custers, R., Dickinson, A., Dweck, C. S., Fishbach, A., Gollwitzer, P. M., Hess, T. M., Isaacowitz, D. M., Mather, M.,… Somerville, L. H. (2014). Mechanisms of motivation-cognition interaction: Challenges and opportunities. Cognitive, Affective & Behavioral Neuroscience, 14(2), 443–472. https://doi.org/10.3758/s13415-014-0300-0

Cabeza, R., Albert, M., Belleville, S., Craik, F. I. M., Duarte, A., Grady, C. L., Lindenberger, U., Nyberg, L., Park, D. C., Reuter-Lorenz, P. A., Rugg, M. D., Steffener, J., & Rajah, M. N. (2018). Maintenance, reserve and compensation: The cognitive neuroscience of healthy ageing. Nature Reviews Neuroscience, 19(11), 701–710. https://doi.org/10.1038/s41583-018-0068-2

Cauvin, S., Moulin, C. J. A., Souchay, C., Kliegel, M., & Schnitzspahn, K. M. (2019). Prospective Memory Predictions in Aging: Increased Overconfidence in Older Adults. Experimental Aging Research, 45(5), 436–459. https://doi.org/10.1080/0361073X.2019.1664471

Chang, L.-H., Shibata, K., Andersen, G. J., Sasaki, Y., & Watanabe, T. (2014). Age-related declines of stability in visual perceptual learning. Current Biology, 24(24), 2926–2929. https://doi.org/10.1016/j.cub.2014.10.041

De Gardelle, V., Le Corre, F., & Mamassian, P. (2016). Confidence as a common currency between vision and audition. PloS One, 11(1), e0147901. https://doi.org/10.1371/journal.pone.0147901

De Gardelle, V., & Mamassian, P. (2014). Does confidence use a common currency across two visual tasks? Psychological Science, 25(6), 1286–1288. https://doi.org/10.1177/0956797614528956

Desender, K., Boldt, A., & Yeung, N. (2018). Subjective Confidence Predicts Information Seeking in Decision Making. Psychological Science, 29(5), 761–778. https://doi.org/10.1177/0956797617744771

Dodson, C. S., Bawa, S., & Krueger, L. E. (2007). Aging, metamemory, and high-confidence errors: A misrecollection account. Psychology and Aging, 22(1), 122–133. https://doi.org/10.1037/0882-7974.22.1.122

Fernandez-Duque, D., Baird, J. A., & Posner, M. I. (2000). Executive attention and metacognitive regulation. Consciousness and Cognition, 9(2 Pt 1), 288–307. https://doi.org/10.1006/ccog.2000.0447

Filippi, R., Ceccolini, A., Periche-Tomas, E., & Bright, P. (2020). Developmental trajectories of metacognitive processing and executive function from childhood to older age. Quarterly Journal of Experimental Psychology, 73(11), 1757–1773. https://doi.org/10.1177/1747021820931096

Flavell, J. H. (1979). Metacognition and cognitive monitoring: A new area of cognitive-developmental inquiry. American Psychologist, 34(10), 906–911. https://doi.org/10.1037/0003-066X.34.10.906

Fleming, S. M., Dolan, R. J., & Frith, C. D. (2012). Metacognition: Computation, biology and function. Philosophical Transactions of the Royal Society of London. Series B, Biological Sciences, 367(1594), 1280–1286. https://doi.org/10.1098/rstb.2012.0021

Fleming, S. M., & Frith, C. D. (2014). Metacognitive Neuroscience: An Introduction. In S. M. Fleming & C. D. Frith (Eds.), The cognitive neuroscience of metacognition (pp. 1–6). Springer. https://doi.org/10.1007/978-3-642-45190-4_1

Fleming, S. M., Huijgen, J., & Dolan, R. J. (2012). Prefrontal contributions to metacognition in perceptual decision making. The Journal of Neuroscience, 32(18), 6117–6125. https://doi.org/10.1523/JNEUROSCI.6489-11.2012

Fu, Y., Yu, S [Shan], Ma, Y., Wang, Y [Yongchang], & Zhou, Y [Yifeng] (2013). Functional degradation of the primary visual cortex during early senescence in rhesus monkeys. Cerebral Cortex, 23(12), 2923–2931. https://doi.org/10.1093/cercor/bhs282

Fuster, J. M. (2000). Executive frontal functions. Experimental Brain Research, 133(1), 66–70. https://doi.org/10.1007/s002210000401

Geurten, M., & Lemaire, P. (2019). Metacognition for strategy selection during arithmetic problem-solving in young and older adults. Neuropsychology, Development, and Cognition. Section B, Aging, Neuropsychology and Cognition, 26(3), 424–446. https://doi.org/10.1080/13825585.2018.1464114

Hansson, P., Rönnlund, M., Juslin, P., & Nilsson, L.-G. (2008). Adult age differences in the realism of confidence judgments: Overconfidence, format dependence, and cognitive predictors. Psychology and Aging, 23(3), 531–544. https://doi.org/10.1037/a0012782

Härting, C., Markowitsch, H.-J., Neufeld, H., Calabrese, P., Deisinger, K., & Kessler, J. (2000). Wechsler Memory Scale, Revised Edition, German Edition. Huber.

Hartshorne, J. K., & Germine, L. T. (2015). When does cognitive functioning peak? The asynchronous rise and fall of different cognitive abilities across the life span. Psychological Science, 26(4), 433–443. https://doi.org/10.1177/0956797614567339

Hasher, L., & Zacks, R. T. (1988). Working memory, comprehension, and aging: A review and a new view. In G. H. Bower (Ed.), Psychology of learning and motivation (Volume 22, pp. 193–225). Academic Press. https://doi.org/10.1016/S0079-7421(08)60041-9

Hertzog, C., & Touron, D. R. (2011). Age differences in memory retrieval shift: Governed by feeling-of-knowing? Psychology and Aging, 26(3), 647–660. https://doi.org/10.1037/a0021875

Huang, J., Gegenfurtner, K. R., Schütz, A. C., & Billino, J. (2017). Age effects on saccadic adaptation: Evidence from different paradigms reveals specific vulnerabilities. Journal of Vision, 17(6), 9. https://doi.org/10.1167/17.6.9

Huang, J., Hegele, M., & Billino, J. (2018). Motivational Modulation of Age-Related Effects on Reaching Adaptation. Frontiers in Psychology, 9, 2285. https://doi.org/10.3389/fpsyg.2018.02285

Kennedy, K. M., Erickson, K. I., Rodrigue, K. M., Voss, M. W., Colcombe, S. J., Kramer, A. F., Acker, J. D., & Raz, N. (2009). Age-related differences in regional brain volumes: A comparison of optimized voxel-based morphometry to manual volumetry. Neurobiology of Aging, 30(10), 1657–1676. https://doi.org/10.1016/j.neurobiolaging.2007.12.020

Kleiner, M. (2010). Visual stimulus timing precision in Psychtoolbox-3: Tests, pitfalls and solutions. Perception, 39, 189.

Kreuzpointner, L., Lukesch, H., & Horn, W. (2013). Leistungsprüfsystem 2, LPS-2: Manual. Hogrefe.

Lachman, J. L., Lachman, R., & Thronesbery, C. (1979). Metamemory through the adult life span. Developmental Psychology, 15(5), 543–551. https://doi.org/10.1037/0012-1649.15.5.543

Lacreuse, A., Raz, N., Schmidtke, D., Hopkins, W. D., & Herndon, J. G. (2020). Age-related decline in executive function as a hallmark of cognitive ageing in primates: An overview of cognitive and neurobiological studies. Philosophical Transactions of the Royal Society of London. Series B, Biological Sciences, 375(1811), 20190618. https://doi.org/10.1098/rstb.2019.0618

Mamassian, P. (2016). Visual Confidence. Annual Review of Vision Science, 2, 459–481. https://doi.org/10.1146/annurev-vision-111815-114630

Maniscalco, B., & Lau, H. (2012). A signal detection theoretic approach for estimating metacognitive sensitivity from confidence ratings. Consciousness and Cognition, 21(1), 422–430. https://doi.org/10.1016/j.concog.2011.09.021

Maniscalco, B., McCurdy, L. Y., Odegaard, B., & Lau, H. (2017). Limited Cognitive Resources Explain a Trade-Off between Perceptual and Metacognitive Vigilance. The Journal of Neuroscience, 37(5), 1213–1224. https://doi.org/10.1523/JNEUROSCI.2271-13.2016

McCurdy, L. Y., Maniscalco, B., Metcalfe, J., Liu, K. Y., Lange, F. P. de, & Lau, H. (2013). Anatomical coupling between distinct metacognitive systems for memory and visual perception. The Journal of Neuroscience, 33(5), 1897–1906. https://doi.org/10.1523/JNEUROSCI.1890-12.2013

Miyake, A., & Friedman, N. P. (2012). The Nature and Organization of Individual Differences in Executive Functions: Four General Conclusions. Current Directions in Psychological Science, 21(1), 8–14. https://doi.org/10.1177/0963721411429458

Monge, Z. A., & Madden, D. J. (2016). Linking cognitive and visual perceptual decline in healthy aging: The information degradation hypothesis. Neuroscience and Biobehavioral Reviews, 69, 166–173. https://doi.org/10.1016/j.neubiorev.2016.07.031

Morales, J., Lau, H., & Fleming, S. M. (2018). Domain-General and Domain-Specific Patterns of Activity Supporting Metacognition in Human Prefrontal Cortex. The Journal of Neuroscience, 38(14), 3534–3546. https://doi.org/10.1523/JNEUROSCI.2360-17.2018

Morgan, M. J., Mason, A. J., & Solomon, J. A. (1997). Blindsight in normal subjects? Nature, 385(6615), 401–402. https://doi.org/10.1038/385401b0

Mueller, S. T., & Piper, B. J. (2014). The Psychology Experiment Building Language (PEBL) and PEBL Test Battery. Journal of Neuroscience Methods, 222, 250–259. https://doi.org/10.1016/j.jneumeth.2013.10.024

Nasreddine, Z. S., Phillips, N. A., Bédirian, V., Charbonneau, S., Whitehead, V., Collin, I., Cummings, J. L., & Chertkow, H. (2005). The Montreal Cognitive Assessment, MoCA: A brief screening tool for mild cognitive impairment. Journal of the American Geriatrics Society, 53(4), 695–699. https://doi.org/10.1111/j.1532-5415.2005.53221.x

O’Callaghan, C., Kveraga, K., Shine, J. M., Adams, R. B., & Bar, M. (2016). Convergent evidence for top-down effects from the “predictive brain”. The Behavioral and Brain Sciences, 39, e254. https://doi.org/10.1017/S0140525X15002599

Owsley, C. (2011). Aging and vision. Vision Research, 51(13), 1610–1622. https://doi.org/10.1016/j.visres.2010.10.020

Owsley, C., Sekuler, R., & Siemsen, D. (1983). Contrast sensitivity throughout adulthood. Vision Research, 23(7), 689–699. https://doi.org/10.1016/0042-6989(83)90210-9

Palmer, E. C., David, A. S., & Fleming, S. M. (2014). Effects of age on metacognitive efficiency. Consciousness and Cognition, 28, 151–160. https://doi.org/10.1016/j.concog.2014.06.007

Pansky, A., Goldsmith, M., Koriat, A., & Pearlman-Avnion, S. (2009). Memory accuracy in old age: Cognitive, metacognitive, and neurocognitive determinants. European Journal of Cognitive Psychology, 21(2-3) 303–329. https://doi.org/10.1080/09541440802281183

Park, D. C., & Reuter-Lorenz, P. (2009). The adaptive brain: Aging and neurocognitive scaffolding. Annual Review of Psychology, 60, 173–196. https://doi.org/10.1146/annurev.psych.59.103006.093656

Peirce, C. S., & Jastrow, J. (1884). On small differences of sensation. Memoirs of the National Academy of Sciences, 3, 73–83.

Perrotin, A., Isingrini, M., Souchay, C., Clarys, D., & Taconnat, L. (2006). Episodic feeling-of-knowing accuracy and cued recall in the elderly: Evidence for double dissociation involving executive functioning and processing speed. Acta Psychologica, 122(1), 58–73. https://doi.org/10.1016/j.actpsy.2005.10.003

Reitan, R. M., & Wolfson, D. (1985). The Halstead--Reitan Neuropsycholgical Test Battery: Therapy and clinical interpretation. Neuropsychological Press.

Roebers, C. M. (2017). Executive function and metacognition: Towards a unifying framework of cognitive self-regulation. Developmental Review, 45, 31–51. https://doi.org/10.1016/j.dr.2017.04.001

Roudaia, E., Farber, L. E., Bennett, P. J., & Sekuler, A. B. (2011). The effects of aging on contour discrimination in clutter. Vision Research, 51(9), 1022–1032. https://doi.org/10.1016/j.visres.2011.02.015

Salthouse, T. A. (1996). The processing-speed theory of adult age differences in cognition. Psychological Review, 103(3), 403–428. https://doi.org/10.1037/0033-295X.103.3.403

Samanez-Larkin, G. R., & Knutson, B. (2015). Decision making in the ageing brain: Changes in affective and motivational circuits. Nature Reviews Neuroscience, 16(5), 278–289. https://doi.org/10.1038/nrn3917

Schütt, H. H., Harmeling, S., Macke, J. H., & Wichmann, F. A. (2016). Painfree and accurate Bayesian estimation of psychometric functions for (potentially) overdispersed data. Vision Research, 122, 105–123. https://doi.org/10.1016/j.visres.2016.02.002

Seidler, R. D., Bernard, J. A., Burutolu, T. B., Fling, B. W., Gordon, M. T., Gwin, J. T., Kwak, Y., & Lipps, D. B. (2010). Motor control and aging: Links to age-related brain structural, functional, and biochemical effects. Neuroscience and Biobehavioral Reviews, 34(5), 721–733. https://doi.org/10.1016/j.neubiorev.2009.10.005

Soderstrom, N. C., McCabe, D. P., & Rhodes, M. G. (2012). Older adults predict more recollective experiences than younger adults. Psychology and Aging, 27(4), 1082–1088. https://doi.org/10.1037/a0029048

Souchay, C., & Isingrini, M. (2004). Age related differences in metacognitive control: Role of executive functioning. Brain and Cognition, 56(1), 89–99. https://doi.org/10.1016/j.bandc.2004.06.002

Stroop, J. R. (1935). Studies of interference in serial verbal reactions. Journal of Experimental Psychology, 18(6), 643–662. https://doi.org/10.1037/h0054651

Toth, J. P., Daniels, K. A., & Solinger, L. A. (2011). What you know can hurt you: Effects of age and prior knowledge on the accuracy of judgments of learning. Psychology and Aging, 26(4), 919–931. https://doi.org/10.1037/a0023379

Valk, S. L., Bernhardt, B. C., Böckler, A., Kanske, P., & Singer, T. (2016). Substrates of metacognition on perception and metacognition on higher-order cognition relate to different subsystems of the mentalizing network. Human Brain Mapping, 37(10), 3388–3399. https://doi.org/10.1002/hbm.23247

Voskuilen, C., Ratcliff, R., & McKoon, G. (2018). Aging and confidence judgments in item recognition. Journal of Experimental Psychology. Learning, Memory, and Cognition, 44(1), 1–23. https://doi.org/10.1037/xlm0000425

Walhovd, K. B., Westlye, L. T., Amlien, I., Espeseth, T., Reinvang, I., Raz, N., Agartz, I., Salat, D. H., Greve, D. N., Fischl, B., Dale, A. M., & Fjell, A. M. (2011). Consistent neuroanatomical age-related volume differences across multiple samples. Neurobiology of Aging, 32(5), 916–932. https://doi.org/10.1016/j.neurobiolaging.2009.05.013

Wechsler, D. (2008). Wechsler Adult Intelligence Scale – Fourth Edition (WAIS–IV). The Psychological Corporation.

West, R. L. (1996). An application of prefrontal cortex function theory to cognitive aging. Psychological Bulletin, 120(2), 272–292. https://doi.org/10.1037/0033-2909.120.2.272

Wong, J. T., Cramer, S. J., & Gallo, D. A. (2012). Age-related reduction of the confidence-accuracy relationship in episodic memory: Effects of recollection quality and retrieval monitoring. Psychology and Aging, 27(4), 1053–1065. https://doi.org/10.1037/a0027686

World Medical Association (2013). Declaration of Helsinki: Ethical principles for medical research involving human subjects. JAMA, 310(20), 2191–2194. https://doi.org/10.1001/jama.2013.281053

Yang, Y., Liang, Z., Li, G., Wang, Y [Yongchang], & Zhou, Y [Yifeng] (2009). Aging affects response variability of V1 and MT neurons in rhesus monkeys. Brain Research, 1274, 21–27. https://doi.org/10.1016/j.brainres.2009.04.015

Yu, S [S.], Wang, Y [Y.], Li, X., Zhou, Y [Y.], & Leventhal, A. G. (2006). Functional degradation of extrastriate visual cortex in senescent rhesus monkeys. Neuroscience, 140(3), 1023–1029. https://doi.org/10.1016/j.neuroscience.2006.01.015

